# Retinoic acid-regulated epigenetic marks identify *Alx1* as a direct target gene required for optic cup formation

**DOI:** 10.1101/2025.06.24.661406

**Authors:** Marie Berenguer, Gregg Duester

**Affiliations:** Development, Aging, and Regeneration Program, Sanford Burnham Prebys Medical Discovery Institute, La Jolla, California, United States of America

**Author notes:** Correspondence should be addressed to G.D.

**Keywords:** Optic cup formation, retinoic acid signaling, RARE, H3K27ac, *Rdh10* knockout, *Alx1*

## Abstract

Retinoic acid (RA) is a transcriptional control agent that regulates several aspects of eye development including invagination of the optic vesicle to form the optic cup, although a target gene for this role has not been previously identified. As loss of RA synthesis in *Rdh10* knockout embryos affects the expression levels of thousands of genes, a different approach is needed to identity genes that are directly regulated by RA. Here, we combined ChIP-seq for epigenetic marks with RNA-seq on eye tissue from wild-type embryos and *Rdh10*-/-embryos that exhibit failure in optic cup formation. We identified a small number of genes with decreased expression when RA is absent that also have decreased presence of a nearby epigenetic gene activation mark (H3K27ac). One such gene was *Alx1* that also has an RA response element (RARE) located near the RA-regulated H3K27ac mark, providing strong evidence that RA directly activates *Alx1*. In situ hybridization studies showed that *Rdh10*-/-embryos exhibit a large decrease in eye *Alx1* expression. CRISPR/Cas9 knockout of *Alx1* resulted in a defect in optic cup formation, thus demonstrating that RA directly activates *Alx1* in order to stimulate this stage of eye development.

**Highlights:**

- The RA requirement for optic cup formation was examined using *Rdh10* knockout embryos.
- Eye RNA-seq and ChIP-seq (H3K27ac) identified *Alx1* as a potential RA target gene.
- *Alx1* exhibits RA-regulated H3K27ac deposition near exon 1 associated with a nearby RARE.
- *Alx1* knockout embryos display a misfolded optic cup with a ventral defect similar to *Rdh10* KO.

## 1. Introduction

Retinoic acid (RA), derived from vitamin A (retinol), controls critical functions during eye development in humans, mice, and zebrafish (Mory et al., 2014; Nedelec et al., 2019; Plaisancie et al., 2016; Slavotinek, 2019; Williams and Bohnsack, 2019). Human studies have associated mutations in four components of the RA signaling pathway (RBP4, STRA6, ALDH1A3, RARB) and two RA target genes (*PITX2, FOXC1*) with anophthalmia or microphthalmia defects that occur during late eye development (Mory et al., 2014; Nedelec et al., 2019; Plaisancie et al., 2016; Slavotinek, 2019; Weisschuh et al., 2006; Williams and Bohnsack, 2019). However, additional RA target genes may exist particulaly for early eye development when optic cup formation occurs from E9.5-E10.5 in mouse embryos. RA controls transcription of key genes by regulating the activity of RA receptors (RARs) that are bound to RA response elements (RAREs) (Cunningham and Duester, 2015; Rhinn and Dolle, 2012). Binding of RA to RARs that are bound to RAREs alters recruitment of nuclear receptor coactivators (NCOAs) known to activate transcription, thus showing that RA activates transcription through RARE enhancers (Ghyselinck and Duester, 2019).

RA is synthesized from retinol in developing optic field tissues by the sequential activities of retinol dehydrogenase-10 (RDH10) (Sandell et al., 2007) and all three retinaldehyde dehydrogenases (ALDH1A1, ALDH1A2, ALDH1A3) (Duester, 2009; Fan et al., 2003; Molotkov et al., 2006). In mouse embryos, *Rdh10* is expressed at E8.75 in optic mesenchyme and at E9.5 in optic vesicle (Sandell et al., 2007). *Aldh1a1* (*Raldh1*) is expressed in the dorsal retina from E9.5-onwards, *Aldh1a2* (*Raldh2*) is expressed in the optic mesenchyme from E8.75-E9.5, and *Aldh1a3* (*Raldh3*) is expressed in the ventral retina from E8.5-onwards (Molotkov et al., 2006). RA is required for folding of the optic vesicle to form the optic cup and ventral retina as shown in *Rdh10*-/-embryos (Sandell et al., 2007) and in *Aldh1a1;Aldh1a2;Aldh1a3* triple knockouts (Molotkov et al., 2006). *Aldh1a1;Aldh1a3* double knockouts form the optic cup (due to *Aldh1a2* RA synthesis), but at later stages they exhibit excessive neural crest-derived perioptic mesenchyme growth and anterior segment defects; in addition, *Pitx2* and *Foxc1* were found to be down-regulated in perioptic mesenchyme (Matt et al., 2005; Molotkov et al., 2006). *Pitx2* and *Foxc1* knockouts exhibit anterior segment eye defects, but as they still generate the optic cup (Evans and Gage, 2005; Kidson et al., 1999), Thus, RA likely regulates at least one additional gene needed for optic cup formation.

We previously provided evidence that *Pitx2* has a nearby RARE and may thus be a direct RA target gene important for late aspects of eye formation (Kumar and Duester, 2010). However, there has been no success in identifying any RA target gene for optic cup formation. Identification of direct RA target genes is difficult as thousands of RAREs are observed in the mouse and human genomes (Lalevee et al., 2011; Moutier et al., 2012) plus the expression of thousands of genes is altered when RA is lost or added (Paschaki et al., 2013; Su and Gudas, 2008). In order to fully understand eye RA signaling, it is essential to identify direct RA target genes that are activated in the developing eye.

As epigenetic studies have shown that histone H3 K27 acetylation (H3K27ac) associates with gene activation (Rada-Iglesias et al., 2011), we propose that genes possessing nearby H3K27ac marks that are decreased by loss of RA may point to direct transcriptional targets of RA. Here, we performed genomic ChIP-seq for H3K27ac and RNA-seq studies on E10 mouse eye tissue from wild-type and *Rdh10*-/-mouse embryos lacking RA synthesis to globally identify direct RA target genes for embryonic eye. Candidate target genes were defined as those with decreased mRNA levels that also have a nearby H3K27ac epigenetic mark that is decreased. This approach identified *Alx1* as a new candidate direct RA target gene, and CRISPR/Cas9 knockout studies demonstrated that *Alx1* is required for normal optic cup formation.

## 2. Methods

### 2.1. Ethics statement

All mouse studies conformed to the regulatory standards adopted by the Institutional Animal Care and Use Committee at the SBP Medical Discovery Institute which approved this study under Animal Welfare Assurance Number A3053-01 (approval #24-080). Animal care and use protocols adhered to the guidelines established by the National Institutes of Health (USA).

### 2.2. Generation of *Rdh10*-/-mouse embryos and isolation of eye tissue

*Rdh10*-/-mice have been previously described (Chatzi et al., 2013). E10 *Rdh10*-/-embryos were generated via timed matings of heterozygous parents; genotyping was performed by PCR analysis of yolk sac DNA. E10 eye tissue from wild-type and *Rdh10*-/-embryos was released from the head of the embryo by dissecting the frontal region containing both optic vesicles and frontonasal region.

### 2.3. RNA-seq analysis

Total RNA was extracted from E10 eye tissue (three pools each of three of wild-type and *Rdh10*-/-eye tissues) and RNA libraries were prepared using the NEB Next Ultra II Directional RNA Library Prep Kit with the PolyA selection module. Sequencing was performed on a NovaSeq platform generating 40 million reads per sample with single read lengths of 75 bp generating 25MM PE50 reads per sample. Sequences were aligned to the mouse mm10 reference genome using TopHat splice-aware aligner; transcript abundance was calculated using Expectation-Maximization approach; fragments per kilobase of transcript per million mapped reads (FPKM) was used for sample normalization; Generalized Linear Model likelihood ratio test in edgeR software was used as a differential test. High throughput DNA sequencing was performed in the La Jolla Immunology Genomics Core. The RNA-seq data is available at GSE297809.

### 2.4. Chromatin immunoprecipitation (ChIP) sample preparation for ChIP-seq

For ChIP-seq we used eye tissue from E10 wild-type or *Rdh10*-/-embryos dissected in modified PBS, i.e. phosphate-buffered saline containing 1X complete protease inhibitors (concentration recommended by use of soluble EDTA-free tablets sold by Roche #11873580001) and 10 mM sodium butyrate as a histone deacetylase inhibitor (Sigma # B5887). Samples were processed similar to previous methods (Berenguer et al., 2020). Dissected eye tissues were briefly centrifuged in 1.5 ml tubes and excess PBS dissection buffer was removed. For cross-linking of chromatin DNA and proteins, 500 μl 1% formaldehyde was added, the eye samples were minced by pipetting up and down with a 200 μl pipette tip and then incubated at room temperature for 15 min. To stop the cross-linking reaction, 55 μl of 1.25 M glycine was added and samples were rocked at room temperature for 5 min. Samples were centrifuged at 5000 rpm for 5 min and the supernatant was carefully removed and discarded. A wash was performed in which 1000 μl of ice-cold modified PBS was added and mixed by vortex followed by centrifugation at 5000 rpm for 5 min and careful removal of supernatant that was discarded. This wash was repeated. Cross-linked eye samples were stored at -80C until enough were collected to proceed, i.e. 60 wild-type and 60 *Rdh10*-/-to perform ChIP-seq with two antibodies in duplicate.

Chromatin was fragmented by sonication. Cross-linked eye samples were pooled, briefly centrifuged, and excess PBS removed. 490 μl lysis buffer (modified PBS containing 1% SDS, 10 mM EDTA, 50 mM Tris-HCl, pH 8.0) was added, mixed by vortexing, then samples were incubated on ice for 10 min. Samples were divided into four sonication microtubes (Covaris AFA Fiber Pre-Slit Snap-Cap 6×16 mm, #520045) with 120 μl per tube. Sonication was performed with a Covaris Sonicator with the following settings - Duty: 5%, Cycle: 200, Intensity: 4, #Cycles: 10, 60 sec each for a total sonication time of 14 min. The contents of the four tubes were re-combined by transfer to a single 1.5 ml microtube which was then centrifuged for 10 min at 13,000 rpm and the supernatants transferred to a fresh 1.5 ml microtube. These conditions were found to shear eye tissue DNA to an average size of 300 bp using a 5 μl sample for Bioanalyzer analysis. At this point 20 μl was removed for each sample (wild-type eye tissue and also *Rdh10*-/-eye tissue) and stored at -20C to serve as input DNA for ChIP-seq.

Each sample was divided into four 100 μl aliquots to perform immunoprecipitation with ChIP-grade antibodies for H3K27ac (Active Motif, Cat#39133) in duplicate. Immunoprecipitation was performed using the Pierce Magnetic ChIP Kit (Thermo Scientific, #26157) following the manufacturer’s instructions. The immunoprecipitated samples and input samples were subjected to reversal of cross-linking by adding water to 500 μl and 20 μl 5 M NaCl, vortexing and incubation at 65C for 4 hr; then addition of 2.6 μl RNase (10 mg/ml), vortexing and incubation at 37C for 30 min; then addition of 10 μl 0.5 M EDTA, 20 μl 1 M Tris-HCl, pH 8.0, 2 μl proteinase K (10 mg/ml), vortexing and incubation at 45C for 1 hr. DNA was extracted using ChIP DNA Clean & Concentrator (Zymo, # D5201 & D5205), After elution from the column in 50 μl of elution buffer, the DNA concentration was determine using 2 μl samples for Bioanalyzer analysis. The two input samples ranged from 16-20 ng/μl and the immunoprecipitated samples ranged from 0.1-0.2ng/μl (5-10 ng per 60 pooled eye tissues). For ChIP-seq, 2 ng was used per sample to prepare libraries for DNA sequencing.

### 2.5. ChIP-seq genomic sequencing and bioinformatic analysis

Libraries for DNA sequencing were prepared according to the instructions accompanying the NEBNext DNA Ultra II kit (catalog # E7645S; New England Biolabs, Inc). Libraries were sequenced on the NextSeq 500 following the manufacturer’s protocols, generating 40 million reads per sample with single read lengths of 75 bp. Adapter remnants of sequencing reads were removed using cutadapt v1.18 (Martin, 2011). ChIP-Seq sequencing reads were aligned using STAR aligner version 2.7 to Mouse genome version 38 (Dobin et al., 2013). Homer v4.10 (Heinz et al., 2010) was used to call peaks from ChIP-Seq samples by comparing the ChIP samples with matching input samples. Homer v4.10 was used to annotate peaks to mouse genes and quantify reads count to peaks. The raw reads count for different peaks were compared using DESeq2 (Love et al., 2014). P values from DESeq2 were corrected using the Benjamini & Hochberg (BH) method for multiple testing errors (Benjamini and Hochberg, 1995). Peaks with BH corrected p value <0.05 (BHP<0.05) were selected as significantly differentially marked peaks. Transcription factor binding sites motif enrichment analyses were performed using Homer v4.10 (Heinz et al., 2010) to analyze the significant RA-regulated ChIP-seq peaks for RARE sequences; DR1 RAREs relate to the TR4(NR),DR1 motif; DR2 RAREs relate to Reverb(NR),DR2; and DR5 RAREs relate to RAR:RXR(NR),DR5. High throughput DNA sequencing was performed in the Sanford Burnham Prebys Genomics Core and bioinformatics analysis was performed in the Sanford Burnham Prebys Bioinformatics Core. The ChIP-seq data is available at GSE298400.

### 2.6. Generation of mutant embryos by CRISPR/Cas9 mutagenesis

CRISPR/Cas9 gene editing was performed using Alt-R CRSIPR/Cas9 tracrRNA (Integrated DNA Technologies, Inc.) combined with crRNAs. Two crRNA guides targeting *Alx1* exon 1 were used to generate frameshift null mutations. crRNAs were designed with maximum specificity using the tool at Integrated DNA Technologies, Inc. to ensure that each crRNA had no more than 17 out of 20 matches with any other site in the mouse genome and that those sites are not located within exons of other genes. The guides used here for targeting the mouse *Alx1* gene were as follows: TTTTACGGCAAAGCGACGGC and AGCATCACGTGCGCCTGGAC. Injection of mouse fertilized eggs and generation of *Alx1* CRISPR embryos was performed using methods previously described by our laboratory (Berenguer et al., 2020; Kumar et al., 2016).

### 2.7. In situ hybridization gene expression analysis

E10.5 embryos were fixed in paraformaldehyde at 4°C overnight, dehydrated into methanol, and stored at -20°C. Detection of mRNA was performed by whole mount in situ hybridization as previously described (Sirbu and Duester, 2006).

### 2.8. Analysis of eye morphology

Histological examination of embryonic eyes was performed on paraffin-sectioned tissues stained with hematoxylin/eosin as previously described (Fan et al., 2003).

## 3. Results

### 3.1. Comparison of H3K27ac ChIP-seq and RNA-seq for *Rdh10*-/-eye tissue

As direct RA target genes for optic cup formation are unknown, we used a combined RNA-seq and ChIP-seq approach that we described previously (Berenguer et al., 2020) to globally identify direct RA target genes for eye development; i.e. genes that when RA is missing exhibit a significant decrease in expression (analyzed by RNA-seq) and also a significant decrease in H3K27ac, an epigenetic mark associated with gene activation (analyzed by ChIP-seq).

Embryonic eye tissues were obtained from E10 embryos by dissected just the frontal region of the head containing both optic vesicles. We performed RNA-seq analysis comparing E10 eye tissue from wild-type embryos and *Rdh10*-/-embryos that lack the ability to produce RA in the entire head including the optic vesicles and surrounding mesenchyme (Chatzi et al., 2013). This analysis identified genes whose mRNA levels in eye tissue are significantly decreased (359 genes) or increased (1040 genes) when RA is absent (FPKM>0.5; a cut-off of log2 <-0.85 or >0.85 was employed; RNA-seq data available at GEO under accession number GSE297809).

We performed ChIP-seq analysis for the H3K27ac epigenetic mark comparing E10 eye tissue from wild-type and *Rdh10*-/- embryos isolated as described above. This analysis identified RA-regulated H3K27ac ChIP-seq peaks that were either decreased (located within or near 3340 genes) or increased (located within or near 3357 genes) using a log2 cut-off of <-0.51 or >0.51; ChIP-seq data is available at GEO under accession number GSE298400.

In order to identify genes that are good candidates for being transcriptionally activated or repressed by RA (RA target genes), we compared our RNA-seq and H3K27ac ChIP-seq results to identify RA-regulated ChIP-seq peaks where nearby genes have significant changes in expression in wild-type vs *Rdh10*-/-based on RNA-seq. We found 81 genes that exhibit both decreased expression and decreased peaks for H3K27ac, plus 202 genes that exhibit increased expression and increased peaks for H3K27ac (Fig. 1).

**Fig. 1.**
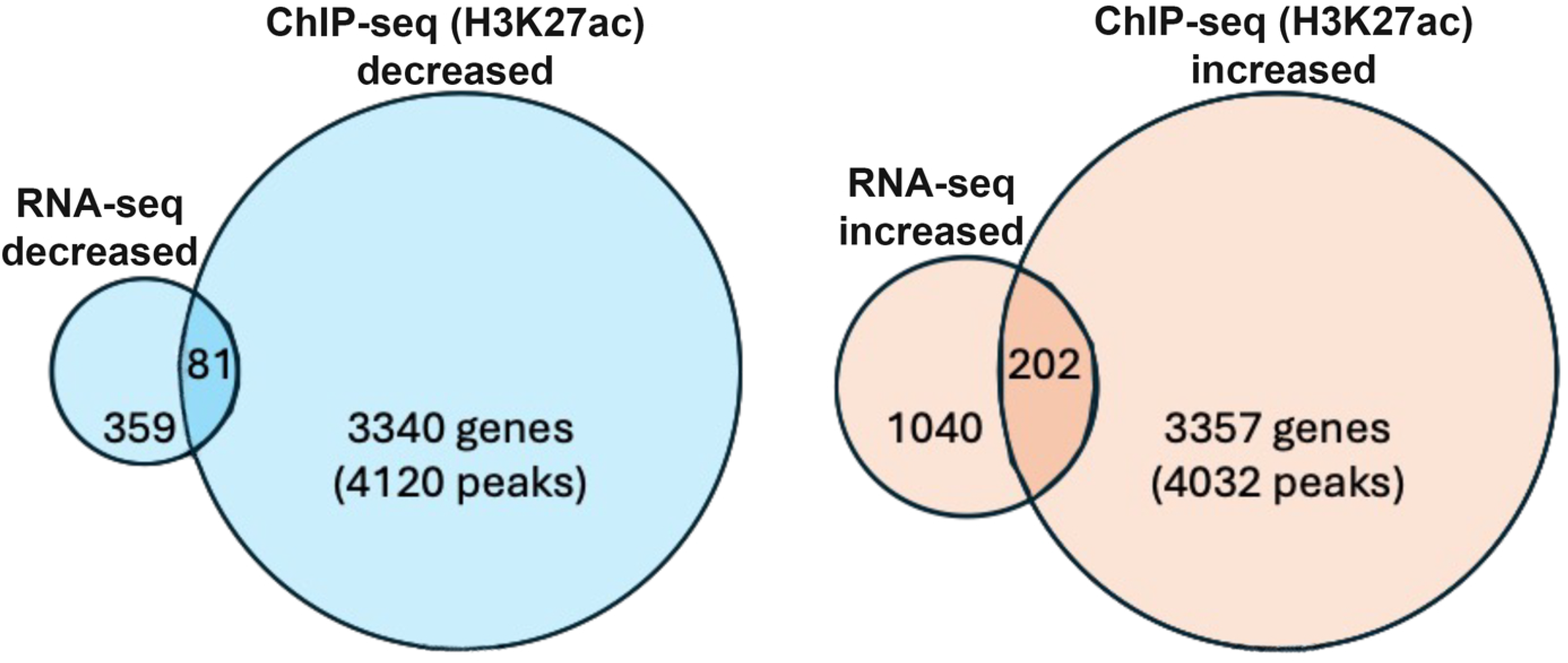
Bioinformatic analysis of genes identified as early eye direct RA target genes. Venn diagram showing the number of genes that have both RA-regulated expression (decreased or increased) and RA-regulated deposition of nearby H3K27ac marks (decreased or increased) following loss of RA. Individual values for H3K27ac (wild-type vs *Rdh10*-/-) can be found in ChIP-seq data deposited at GEO under accession number GSE298400. Individual values for RNA-seq (wild-type vs *Rdh10*-/-) can be found in the data deposited at GEO under accession number GSE297809.

We focused upon the 81 candidate RA target genes that exhibited both decreased expression and decreased H3K27ac deposition. Our combined RNA-seq and H3K27ac ChIP-seq results for this group appear to be high quality it includes several genes that were already known to be part of the optic vesicle RA regulatory pathway; i.e. *Rarb* (nuclear RA receptor), *Rdh10* (first step of RA synthesis -conversion of retinol to retinaldehyde), and *Aldh1a3* (second step of RA synthesis - conversion of retinaldehyde to RA). These three genes are also included within a group of 12 genes with greatest decreases in expression, and here we show their corresponding changes in H3H27ac deposition (Table 1). Among these genes, seven exhibit decreased H3K27ac including *Rarb, Rdh10*, and *Aldh1a3*. This group includes *Alx1* which has a large value for decreased H3K27ac deposition when RA is missing.

**Table 1.** Comparison of datasets for RNA-seq and ChIP-seq (H3K27ac) from E10 eye regions (*Rdh10* KO vs wild-type) to identify genes that have significant reductions in mRNA expression and significant changes in nearby deposition of H3K27ac when RA is missing.

| Gene with altered expression in *Rdh10* KO | log2 fold change in gene expression: RNA-seq for *Rdh10* KO vs WT | nearby H3K27ac ChIP-seq RA-regulated peak for *Rdh10* KO vs WT (mm10 coordinates) | log2 fold change: H3K27ac ChIP-seq for *Rdh10* KO vs WT |
| --- | --- | --- | --- |
| *Adck4* | -0.5 | chr7: 27232464-27233503 | 1.04 |
| *Hspa8* | -0.498 | chr9: 40798968-40802290 | -0.284 |
| *Sdf2l1* | -0.414 | chr16: 17131820-17133225 | 0.052 |
| *Rad51c* | -0.401 | chr11:87403988-87404734 | 0.206 |
| *Car2* | -0.389 | chr14: 70152974-70154100 | -0.236 |
| *Rarb* | -0.37 | chr14: 16554360-16555273 | -0.165 |
| *Aldh1a3* | -0.367 | chr7:66449969-66450726 | -0.165 |
| *Alx1* | -0.363 | chr10:103027606-103029451 | -0.879 |
| *Cln5* | -0.344 | chr14:103069656-103070583 | 0.341 |
| *Rps14* | -0.342 | chr18: 60774247-60775597 | 0.218 |
| *Manf* | -0.342 | chr9:106891507-106892882 | -0.602 |
| *Rdh10* | -0.341 | chr1:16248679-16249396 | -0.395 |

### 3.2 *Alx1* expression in wild-type vs *Rdh10* KO embryos

*Alx1* has previously been shown to be expressed in mouse craniofacial mesenchyme surrounding the optic vesicles and optic cup (Beverdam and Meijlink, 2001). Here, we compared expression of *Alx1* in E10.5 wild-type and *Rdh10*-/-embryos. We found that loss of RA in *Rdh10*-/-embryos results in a very large decrease in *Alx1* expression in the eye region (Fig. 2). These results corroborate the RNA-seq data showing that *Alx1* expression is decreased in the *Rdh10* KO.

**Fig. 2.**
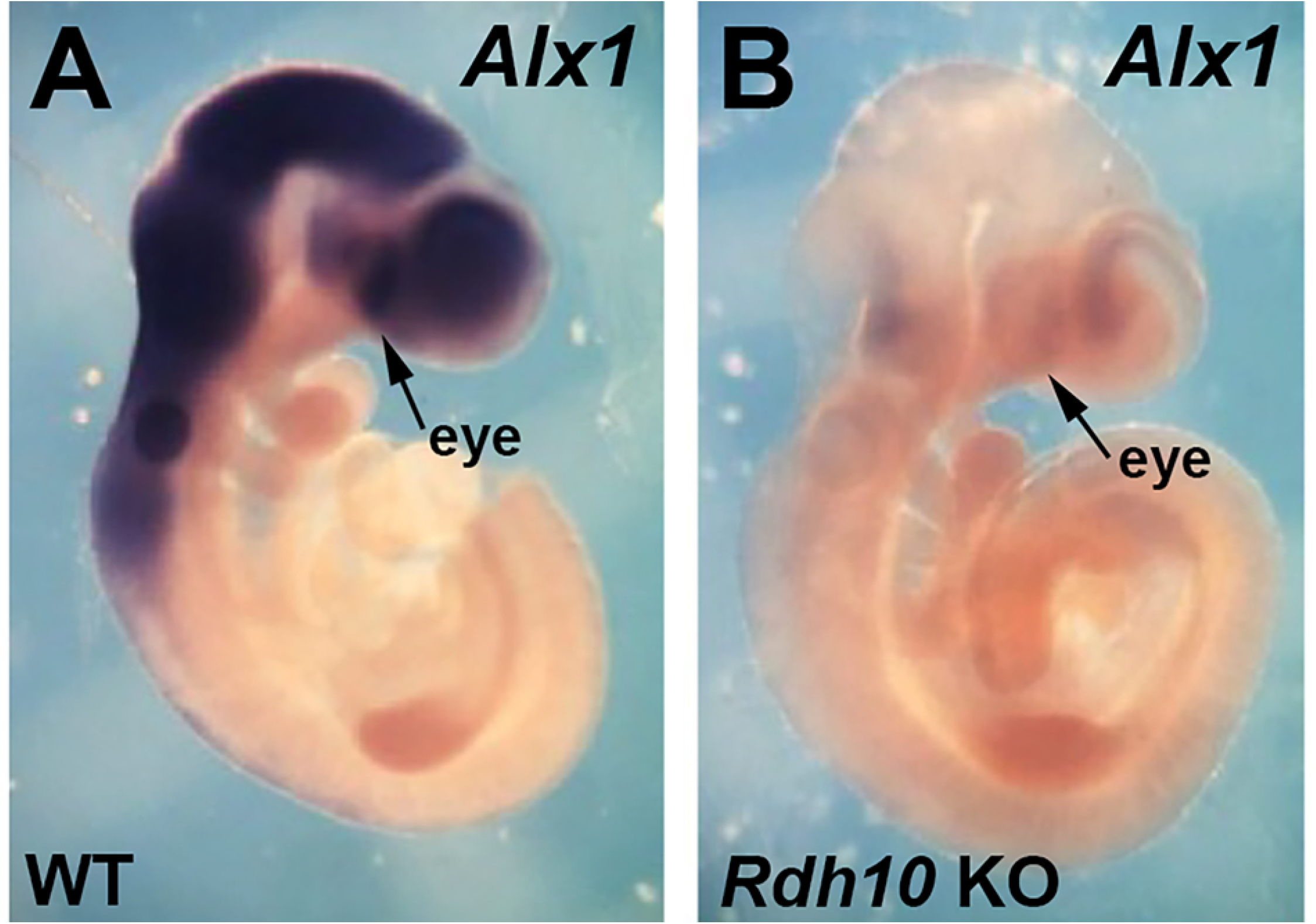
Comparison of *Alx1* expression in E10.5 wild-type vs *Rdh10* KO. (A-B) Shown are mouse E10.5 embryos, either wild-type (WT) or *Rdh10*-/- (*Rdh10* KO), subjected to in situ hybridization to detect *Alx1* mRNA. The arrows point to the eye region where the optic cup is forming.

### 3.3. Identification of a RARE associated with RA-regulated deposition of the H3K27ac epigenetic mark near *Alx1*

A genomic comparison of the H3K27ac ChIP-seq data near the 5’ end of *Alx1* for wild-type and *Rdh10*-/-eye tissue is shown here (Fig. 3). In wild-type, a significant peak of H3K27ac deposition is observed overlapping the transcription start site of *Alx1*, and this peak is reduced in the *Rdh10* KO.

**Fig. 3.**
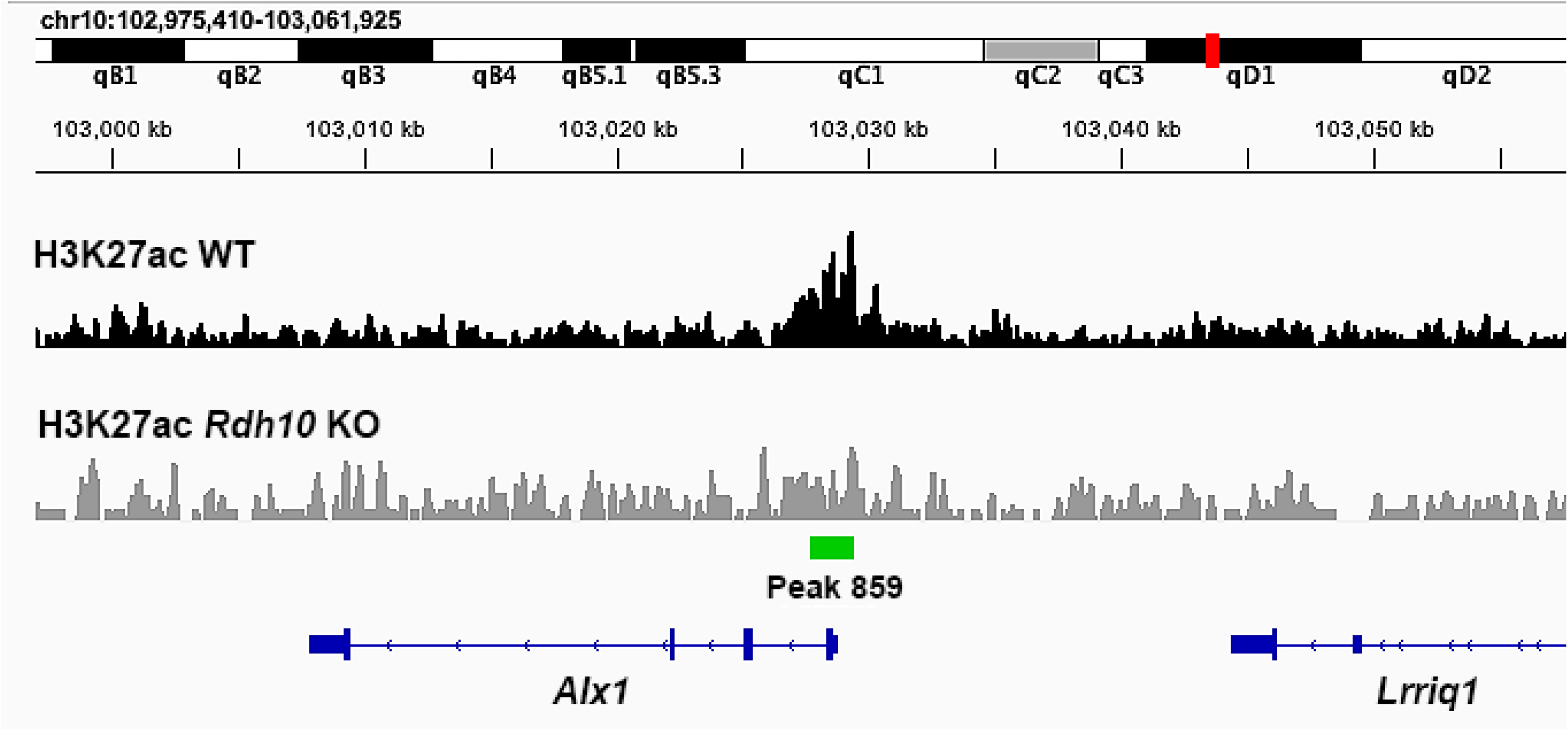
Chromosomal genomic DNA view showing comparison of H3K27ac ChIP-seq data near *Alx1* in wild-type vs *Rdh10*-/-eye tissue. Shown here is the region along mouse chromosome 10 where *Alx1* resides with its transcription direction going from right to left. The ChIP-seq data for H3K27ac from E10.5 eye tissue, either wild-type (WT) or *Rdh10*-/- (*Rdh10* KO), is plotted under this chromosomal region. Observed is peak 859 overlapping the transcription start site for *Alx1* that is significantly decreased in the *Rdh10* K0.

As RA target genes need to be associated with a RARE, the DNA sequences within and near the RA-regulated H3K27ac ChIP-seq peak for *Alx1* were analyzed for the presence of RARE DNA sequences (i.e. direct repeats with spacers of either 1, 2, or 5 bp; DR1, DR2, or DR3) using methods we previously described (Berenguer et al., 2020). We found a DR2 RARE approximately 500 bp upstream of the H3K27ac peak overlapping the *Alx1* transcription start site (Fig. 4). These results provide evidence that *Alx1* is directly regulated by RA during optic cup formation.

**Fig. 4.**
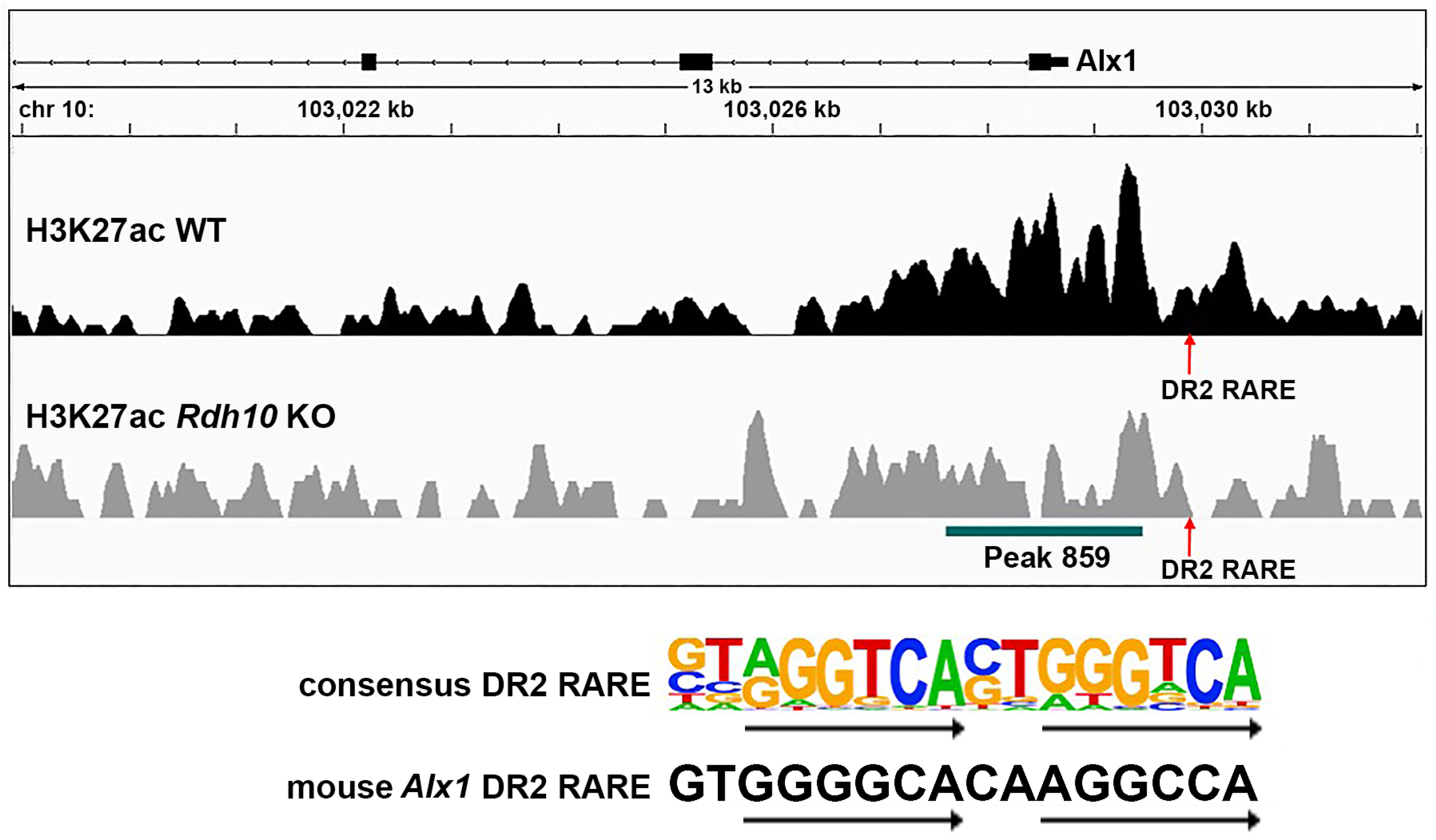
RARE near RA-regulated H3K27ac peak located upstream of the *Alx1* transcription start site. Shown is a mouse chromosome 10 view of the H3K27ac ChIP-seq peaks for wild-type (WT) and *Rdh10*-/-(*Rdh10* KO) from E10.5 eye tissue showing a RA response element (RARE) just upstream of peak 859 which overlaps the *Alx1* transcription start site. This is a DR2 RARE which consists of two 6 bp direct repeats separated by a 2 bp spacer as shown by the consensus DR2 RARE sequence. The DR2 RARE upstream of *Alx1* matches the consensus in 10 of the 12 bp that comprise the two 6 bp repeats, and it matches 1 bp from the 2 bp spacer.

### 3.4. *Alx1* is required for normal optic cup formation

In order to test whether *Alx1* is required for normal optic cup formation, we used CRISPR/Cas9 technology to generate *Alx1* knockout embryos that contain a deletion in exon 1 that introduces a frameshift mutation. Wild-type and *Alx1* CRISPR embryos collected at E10.5 were sectioned in the dorsoventral plane through the eyes. Sections revealed that *Alx1* KO eyes display a failure of the ventral retina to fold properly; *n*=3 (Fig. 5). In the *Alx1* CRISPR embryo, the optic vesicle did not completely fold to generate a normal optic cup with a retinal epithelium that separates from the nearby forebrain epithelium as seen in wild-type. Misfolding of the optic cup may have also contributed to a reduction in the surface ectoderm that will form the cornea and misfolding of the lens vesicle (Fig. 5).

**Fig. 5.**
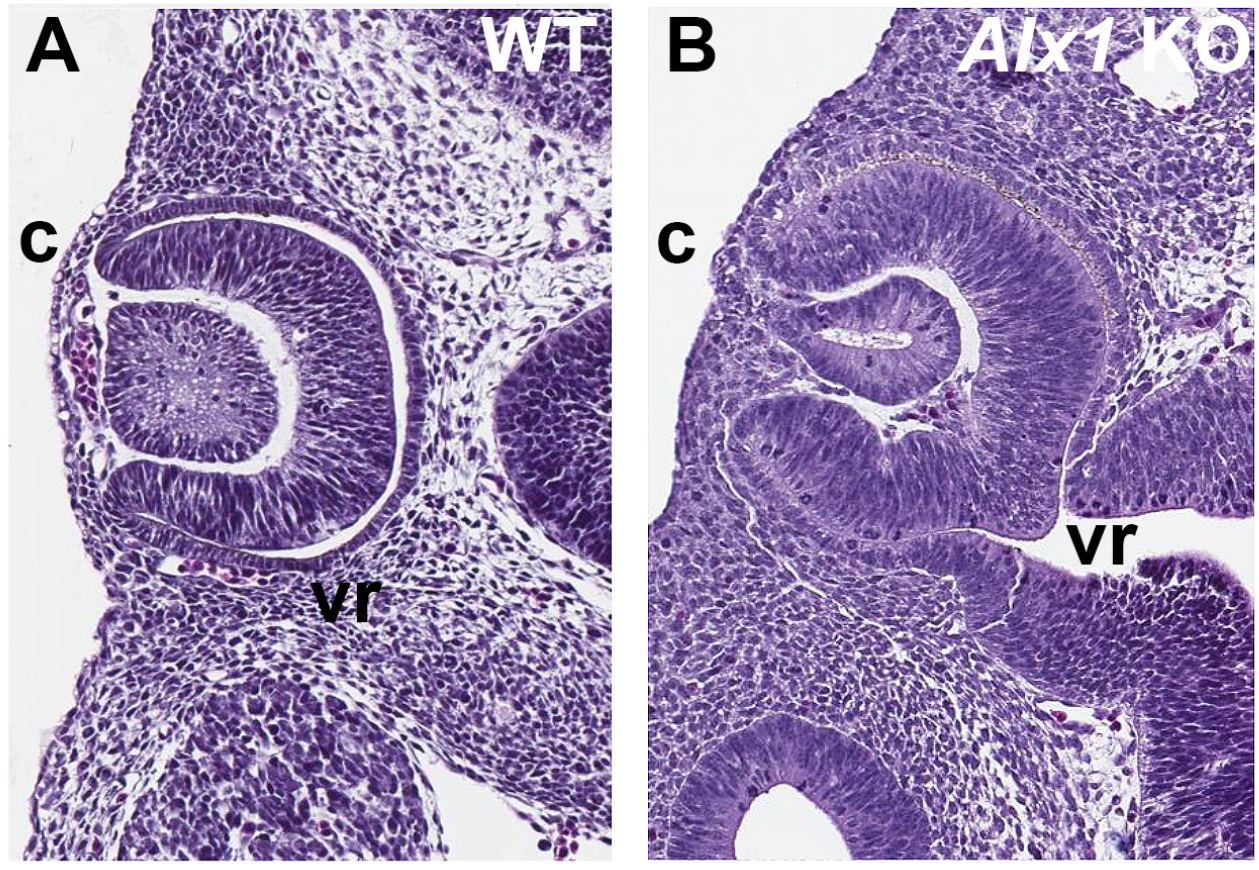
Eye sections comparing wild-type vs *Alx1* CRISPR KO embryos showing optic cup defect. (A-B) Comparison of E10.5 wild-type (WT) and *Alx1* knockout (KO) eyes showing a failure of the ventral retina (vr) to fold properly in the *Alx1* CRISPR embryo and a defect in the appearance of surface ectoderm that will form the cornea (c) as well as a misfolded lens vesicle.

## 4. Discussion

Previous studies have demonstrated that RA signaling is required for folding of the optic vesicle at E9.5 to form the optic cup at E10.5 as well as for further morphogenesis of the optic cup at later stages (Matt et al., 2005; Molotkov et al., 2006; Sandell et al., 2007). However, the identification of an RA target gene required for optic cup formation has not been described. Here, our RNA-seq findings combined with H3K27ac epigenetic ChIP-seq findings comparing wild-type vs *Rdh10*-/-RA-deficient eye tissue has provided a means for identifying new candidate direct RA target genes that may be required for optic cup formation. By focusing on RA-regulated genes that also have a nearby RA-regulated H3K27ac epigenetic mark associated with a RARE, our approach can narrow down the list of RA-regulated genes to those that are direct targets of RA transcriptional control. Our method allows identification of genes that are likely to be transcriptional targets of the RA signaling pathway as opposed to genes whose expression is altered by effects downstream of RA signaling such as changes in expression of other transcription factors or possibly post-transcriptional effects on mRNA abundance that are picked up by RNA-seq analysis.

Here, in our studies on *Rdh10*-/-eye tissue, we were able to narrow down the hundreds of genes identified with RNA-seq that have significant reductions in gene expression to just a few candidate direct RA target genes that also have significant reductions in nearby H3K27ac deposition. From this combined RNA-seq and H3K27ac ChIP-seq comparison, *Alx1* appeared as a very good potential direct RA target gene. This view was further strengthened by analysis of *Alx1* expression in E10.5 embryos showing a large decrease in expression in *Rdh10*-/-compared to wild-type. This view was additionally supported by transcription factor binding site analysis that revealed a DR2 RARE near the RA-regulated H3K27ac peak overlapping the *Alx1* transcription start site. After demonstrating that *Alx1* knockout embryos fail to undergo normal optic cup formation we conclude that *Alx1* is a direct RA target gene whose expression is activated by RA in order to provide instructions for how the optic vesicle folds to form a normal optic cup. As *Alx1* has previously been shown to be expressed in craniofacial mesenchyme surrounding the optic vesicle/cup (Beverdam and Meijlink, 2001), it is possible that *Alx1* controls the migration of cranial neural crest that helps pattern the optic cup similar to how *Pitx2* controls cranial neural crest at later stages of eye morphogenesis (Evans and Gage, 2005; Kidson et al., 1999). Future studies can address the mechanism through which *Alx1* controls optic cup formation.

Our studies repoted here demonstrate the power of combining gene knockout mice, RNA-seq, and ChIP-seq for epigenetic marks to identify direct RA target genes required for a particular developmental process. Here, we have revealed that RA is required to activate expression of *Alx1* that is then required in the process of optic cup formation. This knowledge is essential for understanding the mechanisms underlying how RA signaling controls eye development. This knowledge will also help determine how human eye defects occur, identify genes that may be mutational targets causing human eye defects, and improve strategies to treat eye defects.

## Acknowledgments

We thank the Genomics Core Facility and Bioinformatics Core Facility at SBP Medical Discovery Institute for help with ChIP-seq and RNA-seq analysis. We thank the Animal Resources Core Facility at SBP Medical Discovery Institute for conducting timed-matings to generate mouse embryos.

